# Graphein - a Python Library for Geometric Deep Learning and Network Analysis on Protein Structures and Interaction Networks

**DOI:** 10.1101/2020.07.15.204701

**Authors:** Arian R. Jamasb, Ramon Viñas, Eric J. Ma, Charlie Harris, Kexin Huang, Dominic Hall, Pietro Lió, Tom L. Blundell

## Abstract

Geometric deep learning has well-motivated applications in the context of biology, a domain where relational structure in datasets can be meaningfully leveraged. Currently, efforts in both geometric deep learning and, more broadly, deep learning applied to biomolecular tasks have been hampered by a scarcity of appropriate datasets accessible to domain specialists and machine learning researchers alike. However, there has been little exploration of how to best to integrate and construct geometric representations of these datatypes. To address this, we introduce Graphein as a turn-key tool for transforming raw data from widely-used bioinformatics databases into machine learning-ready datasets in a high-throughput and flexible manner. Graphein is a Python library for constructing graph and surface-mesh representations of protein structures and biological interaction networks for computational analysis. Graphein provides utilities for data retrieval from widely-used bioinformatics databases for structural data, including the Protein Data Bank, the recently-released AlphaFold Structure Database, and for biomolecular interaction networks from STRINGdb, BioGrid, TRRUST and RegNetwork. The library interfaces with popular geometric deep learning libraries: DGL, PyTorch Geometric and PyTorch3D though remains framework agnostic as it is built on top of the PyData ecosystem to enable inter-operability with scientific computing tools and libraries. Graphein is designed to be highly flexible, allowing the user to specify each step of the data preparation, scalable to facilitate working with large protein complexes and interaction graphs, and contains useful pre-processing tools for preparing experimental files. Graphein facilitates network-based, graph-theoretic and topological analyses of structural and interaction datasets in a high-throughput manner. As example workflows, we make available two new protein structure-related datasets, previously unused by the geometric deep learning community. We envision that Graphein will facilitate developments in computational biology, graph representation learning and drug discovery.

**Availability and implementation:** Graphein is written in Python. Source code, example usage and tutorials, datasets, and documentation are made freely available under the MIT License at the following URL: graphein.ai

## 1 Introduction

Geometric deep learning refers to the application of deep learning methods to data with an underlying non-Euclidean structure, such as graphs or manifolds [1]. These methods have already been applied to a number of problems within computational biology and computational structural biology [2, 3, 4, 5, 6, 7, 8], and have shown great promise in the contexts of drug discovery and development [9]. Geometric deep learning libraries have been developed, providing graph representation functionality and in-built datasets - typically with a focus on small molecules [10, 11]. Featurisation schemes and computational analysis of small molecular graphs are a mature area of research. However, data preparation for geometric deep learning in structural biology and interactomics is yet to receive the same attention. Protein function is intricately tied to the underlying molecular structure which is significantly more complex than small molecules. Protein graphs can be populated at different levels of granularity, from atomic-scale graphs similar to small molecules, to residue-level graphs. The relational structure of the data can be captured via spatial relationships or higher-order intramolecular interactions which are not present in small molecule graphs. Furthermore, many biological functions are mediated by interacting biomolecular entities, often through direct physical contacts governed by their 3D structure. As a result, greater control over the data engineering and featurisation process of structural data is required. Little attempt has been made to explore the effect of graph representations of biological structures and to unify structural data and interaction data in the context of machine learning. Graphein is a tool to address these issues by providing flexibility to researchers, decrease the time required for data preparation and facilitate reproducible research.

Proteins form complex three dimensional structures to carry out cellular functions. Decades of structural biology research and recent developments in protein folding have resulted in a large pool of experimentally-determined and modelled protein structures with massive potential to inform future research [12, 13]. However, it is not clear how best to represent these data in machine learning experiments. 3D Convolutional Neural Networks (3DCNNs) have been routinely applied to grid-structured representations of protein structures and sequence-based methods have proved commonplace [14, 15, 16]. Nonetheless, these representations fail to capture relational information in the context of intramolecular contacts and the internal chemistry of the biomolecular structures. Furthermore, these methods are computationally inefficient due to convolving over large regions of empty space, and computational constraints often require the volume of the protein considered to regions of interest, thereby losing global structural information. For instance, in the case of protein-ligand interaction and binding affinity prediction, central tasks in data-driven drug discovery, this often takes the form of restricting the volume to be centred on a binding pocket, thereby losing information about allosteric sites on the protein and possible conformational rearrangements that contribute to molecular recognition. Furthermore, 3D volumetric representations are not translationally and rotationally invariant, deficiencies that are often mitigated using costly data augmentation techniques. Graphs suffer relatively less from these problems as they are translationally and rotationally invariant. Structural descriptors of position can be leveraged and meaningfully exploited by architectures such as Equivariant Neural Networks (ENNs), which ensure geometric transformations applied to their inputs correspond to well-defined transformations of the outputs.

Proteins and biological interaction networks can very naturally be represented as graphs at different levels of granularity. Residue-level graphs represent protein structures as graphs where the nodes consist of amino-acid residues and the edges the relations between them - often based on intramolecular interactions or euclidean distance-based cutoffs. Atom-level graphs represent the protein structure in a manner consistent with small-molecule graph representations, where nodes represent individual atoms and edges the relations between them - often chemical bonds or, again, distance-based cutoffs. The graph structure can further be elaborated by assigning numerical features to corresponding nodes and edges as well as the whole graph. These features can represent, for instance, chemical properties of the residue or atom-type, secondary structure assignments or solvent accessibility metrics of the residue. Edge features can include bond or interaction types, or distances. Graph features can included functional annotations or sequence-based descriptors. In the context of interaction networks, structural data can be overlaid on protein nodes providing a multi-scale view of biological systems and function. Graphein provides a bridge for geometric deep learning into structural interactomics.

Graph representations of proteins have a history of successful applications in machine learning and structural analysis projects in structural biology [17, 18, 19]. Web-servers for computing protein structure graphs exist [20, 21], however the lack of fine-grained control over the construction and featurisation, public APIs for high-throughput programmatic access, ease of integrating data modalities, and incompatibility with deep learning libraries motivated the development of Graphein.

## 2 Related Work

Geometric deep learning methods have demonstrated their suitability for tasks across domains. In part, this has been fuelled by the development of libraries that provide easy access to non-Euclidean data objects and models from the literature. Deep Graph Library (DGL) [10] and PyTorch Geometric [11] are the main open-source frameworks built for PyTorch [22]. Other, less established, tools include: Graph Nets [23] for Sonnet [24]/Tensorflow [25] and Jraph 26] for JAX [27].

In-built dataset support is a common feature of geometric deep learning frameworks. More specialised libraries, such as DGL-LifeSci, DeepChem and TorchDrug, provide datasets, featurisation, neural network layers and pre-trained models for tasks involving small molecules in the life sciences, computational chemistry and drug development [28, 29, 30]. TorchDrug and DeepChem provide reinforcement learning environments to fine tune generative models for physicochemical properties such as drug-likeness (QED) and lipophilicity (LogP). Therapeutics Data Commons provides ML-ready datasets for small molecule and biologics tasks but with no protein structural datasets [31].

Biomolecular tasks are included in many graph representation learning benchmarks. The Open Graph Benchmark (OGB) includes graph property prediction tasks on small molecules, link pre-diction tasks (`ogbl-ppa`) based on protein-protein interaction prediction and a biomedical knowledge graph (`ogbl-biokg`), and a node classification task based on prediction of protein function (`ogbn-proteins`) [32]. The TUDataset contains three biologically-motivated benchmark datasets for graph classification, (`PROTEINS`, `ENZYMES` and `DD`) relevant to applications in structural biology [33]. For `PROTEINS` and `DD` the goal is to predict whether or not a protein is an enzyme and these are derived from the same data under differing graph construction schemes [34, 35]. `ENZYMES` provides a task based on assigning Enzyme Commission (EC) numbers to graph representations of enzyme structures derived from the BRENDA database [36]. More recently, ATOM3D provides a collection of benchmark datasets for structurally-motivated tasks on biomolecules and show leveraging structural information consistently improves performance, and that the choice of architecture significantly impacts performance depending on the context of the task [37].

Whilst tools exist for converting protein structures into graphs, they typically focus on visualisation and leave much to be desired for deep learning practioners. GraProStr is a web-server that enables users to submit structures for conversion into a graph which can be downloaded as textfiles [38]. This provides users with limited control over the construction process, low-throughput and limited featurisation support. Furthermore GraProStr provides no utilities for machine learning or unifying structural and interactomic data o. Mayavi, and GSP4PDB & LIGPLOT provides utilities for visualising protein structures and protein-ligand interaction as graphs, respectively. [39, 40, 41]. Bionoi is a library for representing protein-ligand interactions as voronoi diagram images specifically for applications in machine learning [42],

The protein structure prediction model AlphaFold2 is perhaps the most promising example of geometric deep learning applied to structural biology. Highly-accurate protein structure prediction using AlphaFold2 has been applied at the proteome scale to humans and 20 key model organisms [13, 43]. As a result of these developments, we anticipate a significant amount of growth in the availability of protein structural and interaction data in the coming years. In particular, we identify structural interactomics as an emerging area for geometric deep learning as sparse structural coverage of the interactome can be infilled with modelled structures. The question of how to best leverage and integrate these data with other modalities remains. A recent review of biomedical knowledge graph datasets identifies graph composition, feature and metadata incorporation and reproducibility as key challenges [44]. We have developed Graphein to address these issues and ensure these data are accessible to computational scientists.

## 3 Graphein

Graphein provides utilities for constructing geometric representations of protein and RNA structures, protein-protein interaction networks, and gene regulatory networks. The library is designed for both novice and expert users through the use of a high or low-level API. The high-level API takes standard biological identifiers and a configuration object as input to yield basic geometric representations of the input data. The low-level API offers a detailed customisation of the graph selection from the input data, allowing users to define their own data preparation, graph construction and featurisation functions in a consistent manner. Graphein is built on the PyData Stack to allow for easy inter-operability with standard scientific computing tools and deep learning framework agnosticism. Graphein is organised into submodules for each of the modalities it supports (Figure 2).

**Figure 1:**
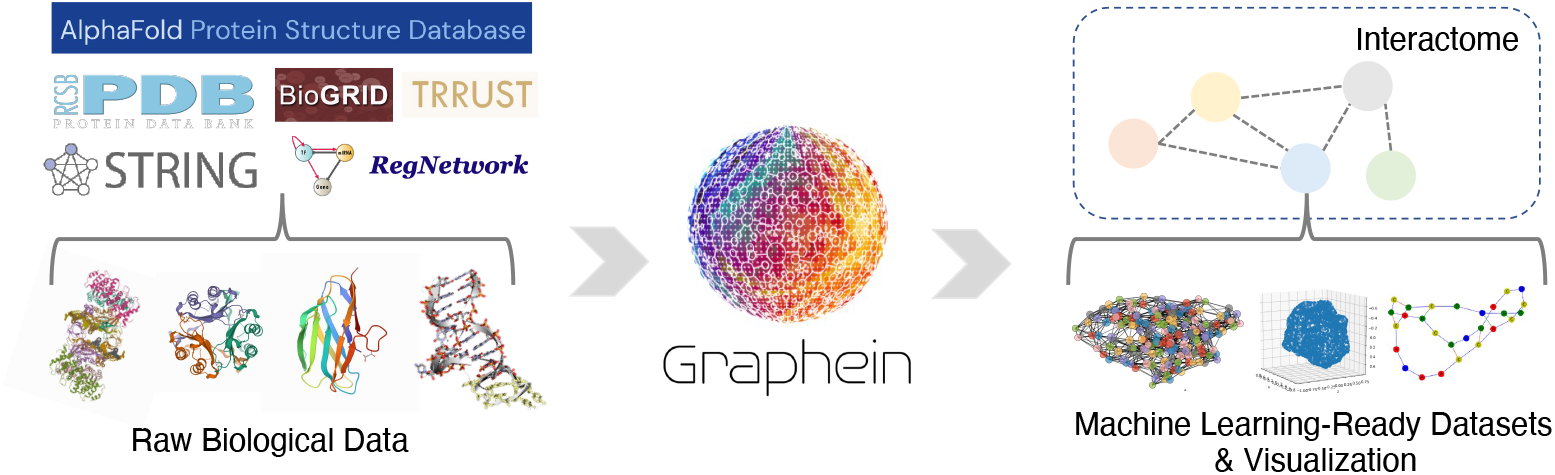
Graphein rapidly transforms and integrates raw biological data into actionable machine learning-ready datasets.

**Figure 2:**
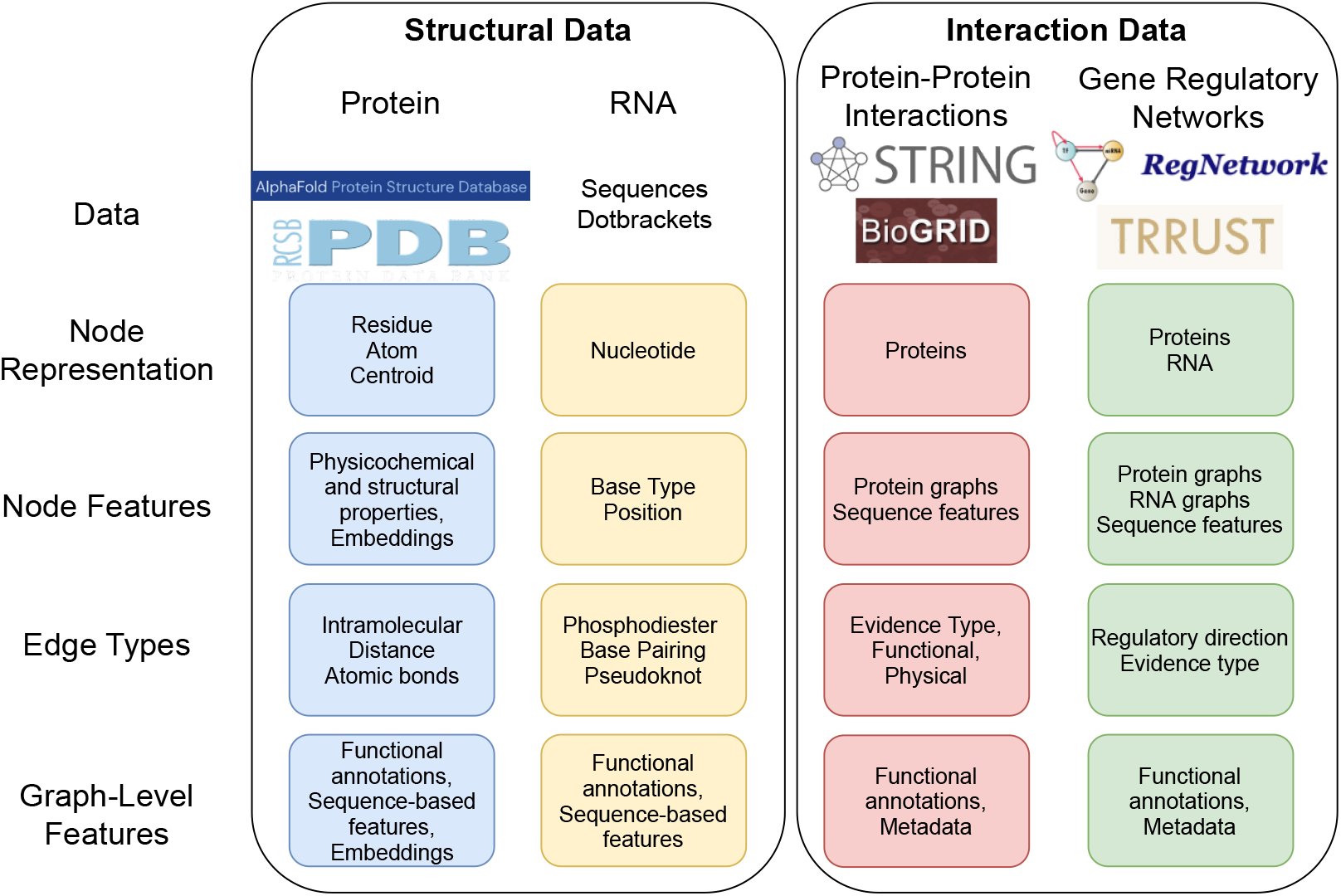
Overview of graph and mesh construction and featurisation schemes for data modalities supported by Graphein. Modules are inter-operable allowing protein or RNA structure graph construction to be applied to nodes in regulatory networks.

### 3.1 Protein Structure Graphs

Graphein interfaces with the PDB and the AlphaFold Structure Database to create geometric representations of protein structures. Furthermore, users can supply their own `.pdb` files, enabling pre-processing with standard bioinformatics tools and pipelines. An overview of featurisation schemes is provided in Supplementary Information A.

#### 3.1.1 Node Representations

Graphs can be constructed for all chains contained within a polypeptide structure, or for a userdefined selection of chains. This is useful in contexts where regions of interest on a protein may be localised to a single chain. For residue-level graphs, users can choose between atom-based positional information, or sidechain centroid. Sidechain centroids are calculated as the centre of gravity of the deprotonated residue. Residue-level graph nodes can be featurised with a one-hot encoding of amino acid type, physicochemical and biochemical properties retrieved from the ExPaSY ProtScale [45] which includes 61 descriptors such as iso-electric points, mutability and transmembrane tendencies. Additional numerical features can be retrieved from AAIndex [46]. Low-dimensional embeddings of amino acid physicochemical properties are provided from Kidera et al. [47] and Meiler et al [48]. In addition to fixed embeddings, sequence embeddings can be retrieved from large pre-trained language models, such as the ESM-1b Transformer model [49] and BioVec [50]. Secondary structural information can be included via a one-hot encoded representation of eight state secondary structure and solvent accessibility metrics (ASA, RSA, SSA) computed by DSSP [51]. *x, y, z* positions are added as node features. Functionality for user-defined node or edge features is also provided with useful utilities allowing for computation or aggregation of features over constituent chains. Figure 2 illustrates an overview of the mesh and graph construction methods as well as the node and edge featurisation schemes; Figure 3 shows example visualisations of graph and meshes produced by Graphein.

**Figure 3:**
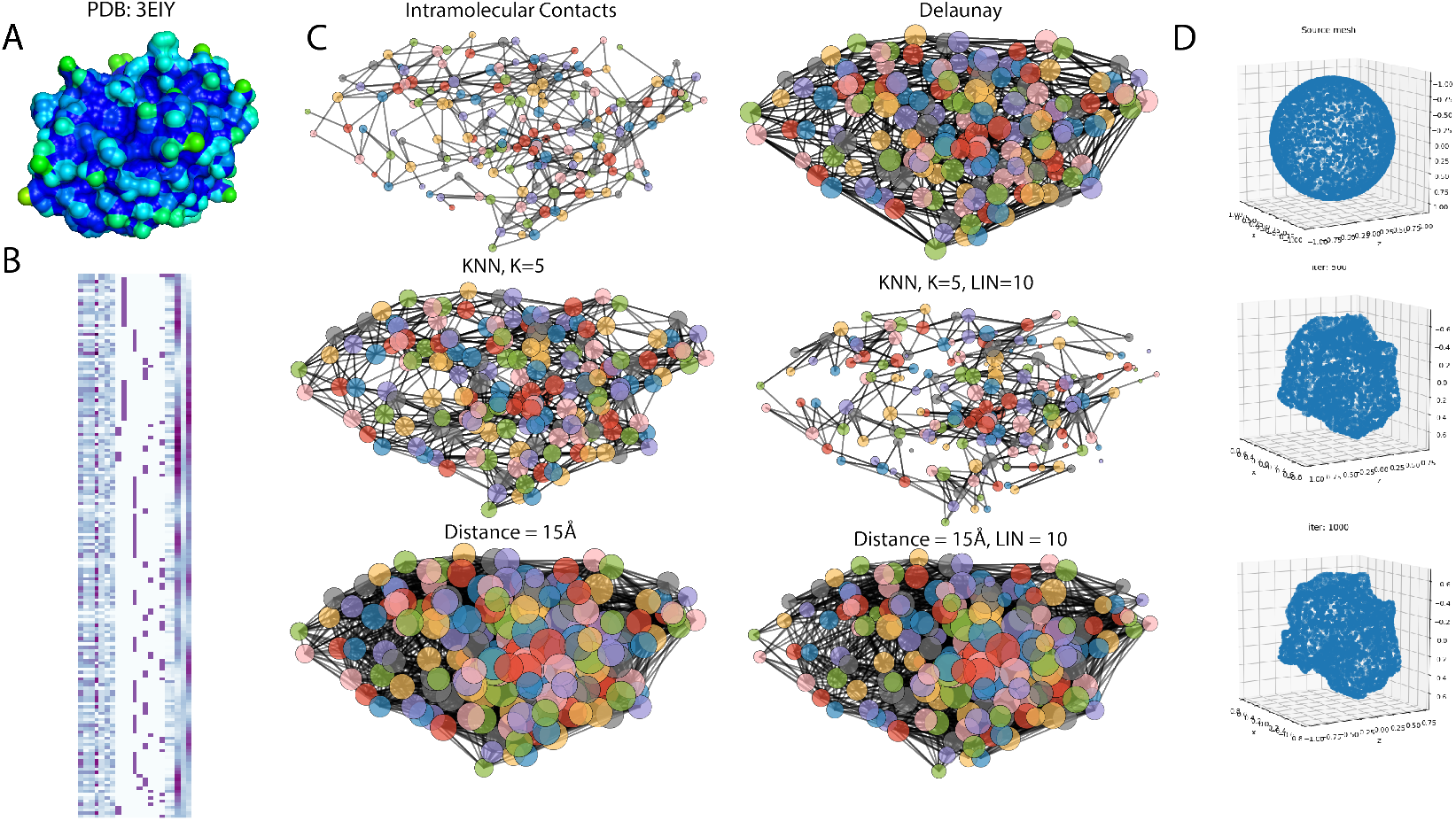
Example protein structure outputs from Graphein. **(A)** Example protein surface (3eiy). **(B)** Example node feature matrix for a residue-level graph. Features include low-dimensional amino acid embeddings, secondary structure assignments, solvent accessibility and *x, y, z* positions of nodes. **(C)** Graphs computed from *α*-carbons in the example protein under a variety of edge construction schemes. Node sizes correspond to degree. **(D)** Deformation of an icosphere to the example protein mesh constructed by Graphein in PyTorch3D

#### 3.1.2 Edge Representations

Graphein provides utilities in the high-level API for a number of edge-construction schemes. The low-level API provides a simple and intuitive way for users to define novel edge construction schemes. Edge construction methods are organised into distance-based, intramolecular interaction-based, and atomic structure-based submodules. Each of these edge construction methods are composable to produce multirelational graphs. This is particularly useful for models that operate on different levels to capture varying aspects of the underlying network. As a motivating example, a multi-track approach has been successfully applied to the protein folding problem [52].

Functionality for computing intramolecular graph edges is provided through distance-based heuristics as well as through an optional dependency, GetContacts [53]. Euclidean distance-based edges can be computed with a user-defined threshold. Functionality for constructing *k*-nearest neighbour graphs, where two vertices are connected by an edge if they are among the *k* nearest neighbours by Euclidean distance is included. Graph edges can also be added on the basis of the Delaunay triangulation. Delaunay triangles correspond to joining points that share a face in the 3D Voronoi diagram of the protein structures. For distance-based edges, a Long Interaction Network (LIN) parameter controls the minimum required separation in the amino acid sequence for edge creation. This can be useful in reducing the number of noisy edges under distance-based edge creation schemes. Edge featurisation for atom-level graphs is provided by annotations of bond type and ring status.

### 3.2 Protein Structure Meshes

Geometric deep learning applied to surface representations of protein structures have demonstrated promise on a variety of tasks in the context of structural biology and structural interactomics [8, 54]. The protein structure mesh module consists of a wrapper for PyMOL, a commonplace molecular informatics visualisation tool, and Pytorch3D [55]. PyMol is used to produce a `.Obj` file from either a PDB accession code or a provided `.PDB` file, enabling the use of pre-processed structures. Pytorch3D is used to produce a tensor-based representation of the protein surface as vertices and faces. Users can pass any desired parameters or commands controlling the surface calculation to PyMol via a configuration object. These parameters include specifying solvent inclusion, solvent probe radius, surface mode (*{triangles, dots}*), surface quality (resolution of mesh). We provide sane defaults for first-time users. To our knowledge, this is the first application of PyTorch3D for protein structure data.

### 3.3 RNA Structures

Ribonucleic Acid (RNA) is a nucleotide biopolymer capable of forming higher-order structural arrangements through self-association mediated by complementary base pairing interactions. Graphein provides utilities for constructing secondary structure graphs of RNA structures, taking as input an RNA sequence and an associated string representation of the secondary structure in dotbracket notation [56]. Graphs can be constructed using two types of bonding between nucleotides: phosphodiester bonds between adjacent bases, and base-pairing interactions between complementary bases specified by the dotbracket string (Figure 4). Graphein also supports addition of pseudoknots - structural motifs composed of interactions between intercalated hairpin loops specified in the dotbracket structure notation.

**Figure 4:**
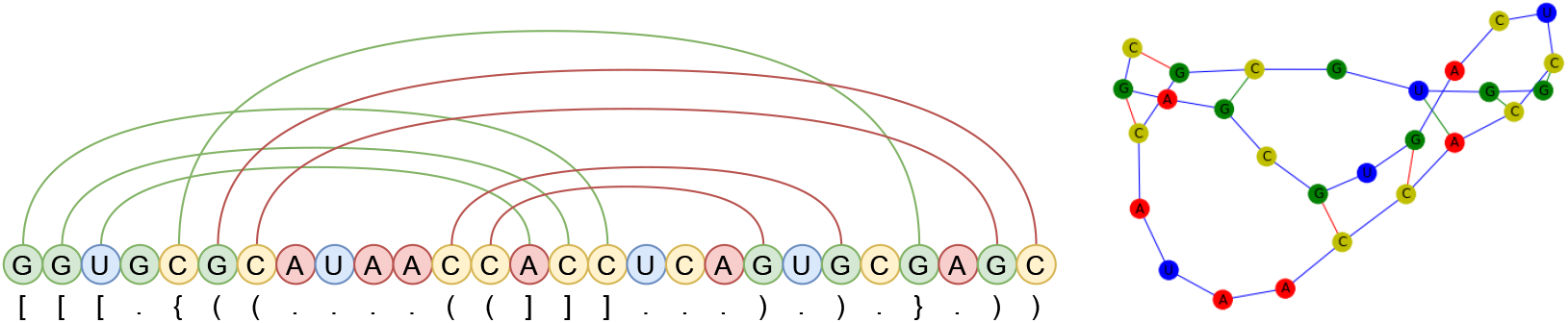
Example RNA Secondary Structure Graph. RNA Secondary structures can be represented as dotbracket strings and multi-relational graphs. Blue edges indicate phosphodiester backbone linkages, red edges indicate base-pairing interactions and green edges indicate pseudoknot pairings.

**Figure 5:**
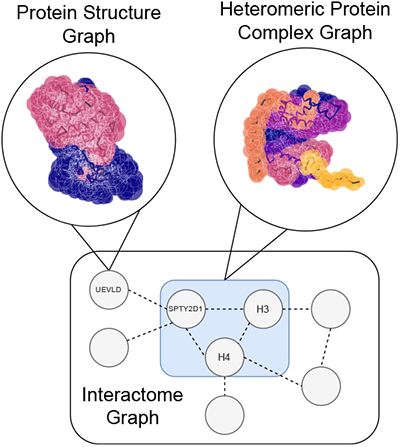
Graphein can facilitate the integration of structural and biomolecular interaction data to enable geometric deep learning research in structural interactomics. 3D visualisations of graphs are generated using Graphein.

### 3.4 Interaction Networks

Interactomics presents a clear application of geometric deep learning as these data are fundamentally relational in structure. Biomolecular entities can be represented as nodes, and their associated functional relationships and physical interactions can be represented as edges with associated metadata, such as the direction and nature of regulation. For a full discussion of applications, datasets and modelling techniques we refer readers to the reviews by [44] and [9]. Graphein implements interaction graph construction from protein-protein interaction and gene regulatory network databases. Interaction graphs integrate networks from several sources and can be constructed in a highly customisable way (see Supplementary Information B, C for a summary of user-definable parameters).

#### 3.4.1 Protein-Protein Interaction Networks

Many of the functional roles of proteins are carried out by larger assemblies of protein complexes and many biological processes are regulated through interactions mediated by physical contacts. Understanding these functions is central to characterising healthy and diseased states of biological systems. Graphein interfaces with widely-used databases of biomolecular interaction data for easy retrieval and construction of graph-based representations of protein-protein interactions.

**STRING** is a database of more than 20 billion known and predicted functional and direct physical protein-protein interactions between 67.6 million proteins across 14,094 organisms [57]. Predicted interactions in STRING are derived from genomic context, high-throughput experimental procedures, conservation of co-expression, text-mining of the literature and aggregation from other databases. STRING is made freely available by the original authors under a Creative Commons BY 4.0 license.

**BioGRID** is a database of 2,127,726 protein and genetic interactions curated from 77,696 publications [58]. BioGRID is made available for academic and commercial use by the original authors under the MIT License.

#### 3.4.2 Gene Regulatory Networks

Gene regulatory networks (GRNs) consist of collections of genes, transcription factors (TFs) and other regulatory elements, and their associated regulatory interactions. Reconstructing transcriptional regulatory networks is a long-standing problem in computational biology in its own right due to its relevance to characterising healthy and diseased states of cells, and these data can provide meaningful signal in other contexts such as multi-modal modelling of biological systems and phenomena. Graphein supports GRN graph construction from two widely-used databases, allowing users to easily unify datasets and construct graph representations of these networks.

**TRRUST** is a database of regulatory interactions for human and mouse interactomes curated from the literature via a sentence-based text-mining approach [59]. The current release contains 8,427 / 6,490 regulatory interactions with associated regulatory directions (activation/repression) over 795 / 827 transcription factors and 2,067 / 1,629 non-transcription factor genes for humans and mice, respectively. TRRUST is made freely available by the original authors for non-commercial research under a Creative Commons Attribution-ShareAlike 4.0 International License.

**RegNetwork** is a database of transcription-factor and miRNA mediated regulatory interactions for humans and mice [60]. RegNetwork is an aggregation of 25 source databases from which the regulatory network is populated and annotated. The latest release contains 14,981 / 94,876 TF-gene, 361 / 129 TF-TF, 21,744 / 25,574 TF-miRNA, 171,477 / 176,512 miRNA-gene and 25,854 / 26,545 miRNA-TF interactions over 1,456 / 1,328 transcription factors, 1,904 / 1,290 miRNAs and 19,719 / 18,120 genes for humans and mice, respectively. The dataset is made publicly available by the original authors.

## 4 Data

### 4.1 Database Interfaces

Graphein interfaces with a number of standard biological data repositories to retrieve data for each modality it supports (summarised in Table 1).

**Table 1:** Graphein database interfaces for data retrieval.

| Name | Description |
| --- | --- |
| **Protein Structure** |  |
| Protein Data Bank (PDB) | Experimentally-determined biomolecular structures |
| AlphaFold Protein Structure Database | Protein structures modelled by AlphaFold2 |
| **Protein-Protein Interaction** |  |
| BioGrid | Protein-protein interactions |
| STRING | Protein-protein interactions |
| **Gene Regulatory Network** |  |
| RegNetwork | TF & miRNA mediated regulatory interactions in humans and mice |
| TRRUST | Regulatory interactions in humans and mice. |

### 4.2 Example Datasets

As example workflows, we make available two graph-based protein structure datasets focussed on tasks where relational inductive biases appear intuitively useful and demonstrate how Graphein can help formulate different tasks from the same underlying dataset.

#### PPISP - Protein Protein Interaction Site Prediction

The first, based on the collections outlined in [61], consists of 420 protein structures, with node labels indicating whether a residue is involved in a protein-protein interaction - a task central to structural interactomics [62]. The data originate from co-crystallised structures of the complexes in the RCSB PDB. The authors make available a set of additional node features based on Position-Specific Scoring Matrices (PSSMs), providing evolutionary context as to which protein-protein interaction sites are typically conserved, which can be incorporated with the structural node features calculated by Graphein. This dataset was used in [63] in conjunction with Graphein to compute the protein structure graph inputs to a Message-Passing Neural Process model which achieved state-of-the-art performance.

#### PSCDB - Protein Structural Change Database

The second dataset, based on Protein Structural Change Database (PSCDB) [64], consists of 904 paired examples of bound and unbound protein structures that undergo 7 classes of conformational rearrangement motion. Prediction of conformational rearrangement upon ligand binding is a longstanding problem in computational structural biology and has significant implications for drug discovery and development. Two tasks can be formulated with this dataset. The first is the graph classification task of predicting the type of motion a protein undergoes upon ligand binding, the second is framing prediction of the rearrangement itself as an edge prediction task between the paired bound and unbound protein structure graphs. These tasks provide utility in improving understanding of protein structural dynamics in drug development, where molecules are typically docked into largely rigid structures with limited flexibility in the binding pockets in high-throughput *in silico* screens. PSCDB is made publicly available by the original authors and we provide a processed version in our repository.

#### ccPDB

We derive four datasets, each with a graph and a node classification task from the ccPDB [65]. The ccPDB provides collections of protein structures and annotations of interactions with various molecular species. The proteins are high-quality, non-redundant sets (25% sequence identity) with maximum resolution of 3 Å, minimum sequence length of 80 residues. Node-level annotations of interaction are provided in each case with the cutoff set at 4 Å. ccPDB is made freely available online.

`PROTEINS_METAL` contains protein structures that bind 7 types of metal ions (Fe, Mg, Ca, Mn, Zn, Co, Ni; *n* = 215 / 1,908 / 1,402 / 521 / 1,660 / 201 / 355).

- `PROTEINS_NUCLEOTIDES` contains protein structures that bind 8 species of nucleotides (ATP, ADP, GTP, GDP, NAD, FAD, FMN, UDP; *n* = 313 / 353 / 83 / 120 / 140 / 172 / 117 / 68)
- `PROTEINS_NUCLEIC` contains protein structures that bind DNA or RNA polymers (*n* = 560 / 415).
- `PROTEINS_LIGAND` contains protein structures that bind 7 species of ligands (SO_4_, PO_4_, NAG, HEM, BME, EDO, PLP; *n* = 3312 / 1299 / 727 / 176 / 191 / 1507 / 65).

## 5 Benchmarks

To demonstrate the ease-of-use of Graphein we apply a selection of geometric deep learning models to protein structure graphs generated by Graphein to these datasets. In particular, we consider two graph construction schemes from the same dataset: one based on hydrogen and peptide bonds (Bond), and another based on K-Nearest Neighbours clustering of the residues (KNN). Full details of our experimental procedure are provided in Supplementary Information D. We consider a graph classification task on `PSCDB`, classifying structures on the basis of the type of structural rearrangement they undergo upon ligand binding. Our results show strong differences in performance between the two schemes and that these differences are architecture-dependent (Table 2). We suggest that this motivates further exploration of the role of graph construction in the context of structural biological data.

**Table 2:** Baseline results for `PSCDB`. The task is graph classification to predict the class of structural rearrangement a given protein undergoes upon ligand binding.

| Edge Type | Bond |  | KNN |  |
| --- | --- | --- | --- | --- |
| Model | Multi-ACC | Macro-F1 | Multi-ACC | Macro-F1 |
| GCN | 0.221±0.009 | 0.110±0.062 | 0.261±0.111 | 0.154±0.084 |
| GraphSAGE | 0.170±0.091 | 0.078±0.046 | 0.247±0.133 | 0.136±0.090 |
| GAT | 0.241±0.066 | 0.145±0.037 | 0.258±0.136 | 0.154±0.105 |

## 6 Machine Learning Utilities

### Conversion

Convenience utilities for converting between NetworkX [66] graph objects and commonly-used geometric deep learning library data objects are provided for DGL and PyTorch Geometric. Underlying graph objects are based on NetworkX, enabling conversion to other formats.

### Adjacency Tensors & Diffusion Matrices

Graphein provides utilities for computation of diffusion matrices (and related adjacency matrices) to (1) facilitate exploration of biological data with models that leverage these representations, and (2) aid in the construction of diffusion matrices for graph neural networks.

### Visualisation

Built-in tools are provided for each of the modalities supported to allow inspection of data in pre and post-processing. Interactive visualisation tools are provided for protein structures.

## 7 Usage

Example usage and workflows are provided in the documentation at: www.graphein.ai. Examples and tutorials are provided as runnable notebooks detailing use of the high and low-level APIs for the data modalities currently supported by Graphein, and the ease of ingesting novel structural datasets into a suite of geometric deep learning benchmarks (see section 5). Source code is made available via GitHub: www.github.com/a-r-j/graphein.

## 8 Conclusion

Geometric deep learning has shown promise in computational biology and structural biology. However, the availability of processed datasets is a research bottleneck. Graphein is a Python library designed to facilitate construction of datasets for geometric deep learning applied to biomolecular structures and interactions. By providing tools for these modalities, we hope to facilitate research in data-driven structural interactomics. In addition, we make available two datasets for protein-protein interaction site prediction (node classification) and protein conformational rearrangement prediction (graph classification and edge prediction).

A current limitation of the library is the lack of support for some informative features based on evolutionary information. For example, the `PPISP` dataset provides PSSMs and the protein folding model, AlphaFold2 is heavily reliant on Multiple Sequence Alignments presenting clear utility for the addition of these features. Whilst graphs are a natural representation of biological interaction data, hypergraphs may provide a higher-fidelity representation of the underlying biological relationships. Many interactions are contextual, which can be represented by hyperedges between several entities required for a functional or structural relationship. We are also interested in addressing representations of dynamics, both in structural data and in interactions as these are central biological components that are beyond the scope of the initial release. These features will be included in subsequent releases and the API design of Graphein makes it simple for users to write and contribute their own workflows. Graphein implements a high-level and low-level API to enable rapid and fine-grained control of data preparation. Graphein is provided as Free Open Source Software under a permissive MIT License which we hope will encourage the community to contribute customised workflows to the library. We hope that Graphein serves to further progress in the field and reduce friction in processing structural and interaction data for geometric deep learning. The library also provides utility in preparing protein structure and interactomics graphs for graph-theoretic and topological data analyses.

## Acknowledgments and Disclosure of Funding

We thank Ben Day, Paul Scherer and Cristian Bodnar for useful feedback on the manuscript and project. We thank Sean Aubin for contributing the interactive visualisation code for protein structure graphs. We thank our users for bug reports, feature suggestions and their support. ARJ is funded by a BBSRC DTP studentship. DH has received funding from both the Wellcome Trust and the United Kingdom Medical Research Council (MRC). TLB thanks the Wellcome Trust for an Investigator Award (200814/Z/16/Z; 2016 -) for support of this research.

## Checklist

1. For all authors…
  a. Do the main claims made in the abstract and introduction accurately reflect the paper’s contributions and scope? [Yes]
  b. Did you describe the limitations of your work? [Yes] **Yes, please see the conclusion (Section 8)**.
  c. Did you discuss any potential negative societal impacts of your work? [Yes] **Please see Supplementary Information E**
  d. Have you read the ethics review guidelines and ensured that your paper conforms to them? [Yes] **We have read the ethics review guidelines**.
2. If you are including theoretical results…
  a. Did you state the full set of assumptions of all theoretical results? [N/A] **We are not including theoretical results**.
  b. Did you include complete proofs of all theoretical results? [N/A] **We are not including theoretical results**.
3. If you ran experiments (e.g. for benchmarks)…
  a. Did you include the code, data, and instructions needed to reproduce the main experimental results (either in the supplemental material or as a URL)? [Yes] **Please see the repository:** https://www.github.com/a-r-j/graphein **and documentation:** www.graphein.ai
  b. Did you specify all the training details (e.g., data splits, hyperparameters, how they were chosen)? [Yes] **A full description is given in Supplementary Information D**
  c. Did you report error bars (e.g., with respect to the random seed after running experiments multiple times)? [Yes] **Please see Table 2**.
  d. Did you include the total amount of compute and the type of resources used (e.g., type of GPUs, internal cluster, or cloud provider)? [Yes] **Please see Supplementary Information D**.
4. If you are using existing assets (e.g., code, data, models) or curating/releasing new assets…
  a. If your work uses existing assets, did you cite the creators? [Yes] **We have cited the appropriate creators in the text**.
  b. Did you mention the license of the assets? [Yes] **We have described the licensing conditions in the text**.
  c. Did you include any new assets either in the supplemental material or as a URL? [Yes] **Please see our repository:** https://www.github.com/a-r-j/graphein **and documentation:** graphein.ai
  d. Did you discuss whether and how consent was obtained from people whose data you’re using/curating? [Yes] **We use publicly available data and specify the original licenses**.
  e. Did you discuss whether the data you are using/curating contains personally identifiable information or offensive content? [N/A] **No datasets include personally identifiable information or offensive content**.
5. If you used crowdsourcing or conducted research with human subjects…
  a. Did you include the full text of instructions given to participants and screenshots, if applicable? [N/A] **We do not use crowdsourcing or human subjects**.
  b. Did you describe any potential participant risks, with links to Institutional Review Board (IRB) approvals, if applicable? [N/A] **We do not use crowdsourcing or human subjects**.
  c. Did you include the estimated hourly wage paid to participants and the total amount spent on participant compensation? [N/A] **We do not use crowdsourcing or human subjects**.

