## Supplementary material for "Graphein - a Python Library for Geometric Deep Learning and Network Analysis on Protein Structures and Interaction Networks": graphein_neurips_supplmentary_revised.pdf

### Graphein - Supplementary material

[1] Department of Biochemistry, University of Cambridge
[2] Department of Computer Science & Technology, University of Cambridge
[3] PyMC Labs
[4] Department of Life Sciences, Imperial College London
[5] Department of Computer Science, Stanford University

Preprint. Under review.

**E Societal Impact** **7**

### A Featurisation Schemes for Protein Structure Graphs

Table 1: Geometric representation options for a protein structure

| *Node Features* | | |
|---|---|---|
| Node Type | Feature | Source |
| **Residue** | | |
| | Molecular Weight | |
| | $x, y, z$ Co-ordinates | |
| | $\phi$ Torsion Angle | DSSP [1] |
| | $\psi$ Torsion Angle | DSSP [1] |
| | Secondary Structure | DSSP [1] |
| | Solvent Accessibility | DDSP [1] |
| | Low-dimensional embeddings of physicochemical properties | Meiler et al. [2] |
| | ExPaSy Protein Scale | [3] |
| | AAIndex Descriptors (various) | [4] |
| | ESM Transformer Protein Language Model Embedding | ESM [5] |
| | BioVec Protein Language Model Embedding | ProtVec [6] |
| **Atom** | | |
| | Atomic Weight | |
| | Covalent Radius | [7] |

| *Edge Types* | | | |
|---|---|---|---|
| Node Type | Edge Type | Features | Source |
| **Residue** | | | |
| | Hydrophobic Interactions | | |
| | Disulpfide Interactions | | |
| | Hydrogen Bonds | | |
| | Ionic Interactions | | |
| | Aromatic Interactions | | |
| | Aromatic-Sulphur Interactions | | |
| | Cation-$\pi$ Interactions | | |
| | Peptide Bonds | | |
| | $\pi$ Stacking Interactions | | [8] |
| | Salt Bridge | | [8] |
| | t Stacking | | [8] |
| | Van der Waals | | [8] |
| **Atom** | | | |
| | | Bond Order | [9] |
| | | Ring Status | |
| **Any** | | | |
| | K-Nearest Neighbours | | |
| | Delaunay Triangulation | | |
| | Distance Threshold | | |

| *Graph-level Features* | | |
|---|---|---|
| | Features | Source |
| **Sequence** | | |
| | Molecular Weight | |
| | ESM Transformer Protein Language Model Embedding | ESM [5] |
| | BioVec Protein Language Model Embedding | ProtVec [6] |
| | Amino Acid Composition | ProPy [10] |
| | Dipeptide Composition | ProPy [10] |
| | Tripeptide Composition | ProPy [10] |
| | Moreau-Broto Autocorrelation | ProPy [10] |
| | Moran Autocorrelation | ProPy [10] |
| | Geary Autocorrelation | ProPy [10] |
| | Sequence-order-coupling Number | ProPy [10] |
| | Quasi-Sequence-Order Descriptors | ProPy [10] |
| | CTD Descriptors | ProPy [10] |

#### B Parameters for protein-protein interaction graphs

##### B.1 General parameters

The following table shows the generic Graphein parameters for protein-protein interaction graphs.

| Parameter | Default | Valid values | Description |
|---|---|---|---|
| protein_list | – | protein IDs (list of string) | Proteins to include in the graph. |
| ncbi_taxon_id | 9606 (human) | integer | NCBI taxon identifier. |
| sources | all sources | 'biogrid', 'string' | List of sources (databases) to retrieve the data from. |
| paginate | True | True, False | Whether to paginate the API calls for the sources that require it. |

##### B.2 BioGRID

The following table shows the Graphein parameters for BioGRID. See also the BioGRID API.

| Parameter | Default | Valid values | Description |
|---|---|---|---|
| searchNames | True | True, False | If True, the interactor OFFICIAL_SYMBOL will be examined for a match with the protein list. |
| searchIds | True | True, False | If True, the interactor ENTREZ_GENE, ORDERED LOCUS and SYSTEMATIC_NAME (orf) will be examined for a match with the protein list. |
| searchSynonyms | True | True, False | If True, the interactor SYNONYMS will be examined for a match with the protein list. |
| searchBiogridIds | True | True, False | If True, the entries in the protein list will be compared to BIOGRID internal IDS which are provided in all Tab2 formatted files. |
| additionalIdentifierTypes | empty | string | Identifier types on this list are examined for a match with the protein list. |
| max | 10000 | integer | Number of results to fetch. Used for pagination. |

| | | | |
|---|---|---|---|
| interSpeciesExcluded | True | True, False | If True, interactions with interactors from different species will be excluded. |
| selfInteractionsExcluded | False | True, False | If True, interactions with one interactor will be excluded. |
| evidenceList | empty | Pipe-separated list of evidence codes from here | Any interaction evidence with its Experimental System in the list will be excluded from the results unless includeEvidence is set to true. |
| includeEvidence | False | True, False | If set to true, any interaction evidence with its Experimental System in the evidenceList will be included in the result |
| excludeGenes | False | True, False | If true, interactions containing genes in the input list will be excluded from the results. |
| includeInteractors | True | True, False | If true, in addition to interactions between genes on the input list, interactions will also be fetched which have only one interactor on the input list. |
| includeInteractorInteractions | False | True, False | If true, interactions between the input list's first order interactors will be included. |
| pubmedList | empty | string | Interactions will be fetched whose Pubmed Id is/ is not in this list, depending on the value of excludePubmeds. |
| excludePubmeds | False | True, False | If False, interactions with Pubmed ID in pubmedList will be included in the results; if 'true' they will be excluded. |
| htpThreshold | 20 | integer | Interactions whose Pubmed ID has more than this number of interactions will be excluded from the results. Ignored if excludePubmeds is False. |

| | | | |
|---|---|---|---|
| throughputTag | 'any' | 'low', 'high', 'any' | If set to 'low or 'high', only interactions with 'Low throughput' or 'High throughput' in the 'throughput' field will be returned. |

##### B.3 STRING

The following table shows the Graphein parameters for STRING. See also the STRING API.

| Parameter | Default | Valid values | Description |
|---|---|---|---|
| network_type | 'functional' | 'functional', 'physical' | Network type: functional (default), physical. |
| add_nodes | 0 | integer | Adds a number of proteins to the network based on their confidence score, e.g., extends the interaction neighborhood of selected proteins to desired value. |
| show_query_node_labels | False | True or False | When available use submitted names in the preferredName column. |

#### C Parameters for gene regulatory networks

##### C.1 General parameters

We download gene regulatory networks from TRRUST and RegNetwork. We build a directed graph where nodes are genes and attributed edges represent regulatory effects (activation, repression, or unknown).

| Parameter | Default | Valid values | Description |
|---|---|---|---|
| gene_list | – | gene symbols (list of string) | Genes to include in the graph. |

#### D Baseline Experiment Details

##### D.1 Data Preparation

Residue-level protein structure graphs are prepared using default Graphein pre-processing: $\alpha$-carbon atoms are used as nodes; heteroatoms are removed from the structure; insertions are removed from the structure, waters are removed from the structure. For the K-NN graphs, we use $k = 5$ and $long_interaction = 0$. For the intramolecular bond graphs we use Hydrogen and backbone peptide bonds only. The only node features considered are $x, y, z$ co-ordinates.

##### D.2 Models

We use three baseline graph representation learning models GCN [11], GraphSAGE [12], and GAT [13], using Pytorch Geometric and Pytorch Lightning for training. For each model, we conduct a hyperparameter tuning sweep using bayesian search on the validation loss. For GCN and GraphSAGE, the hyperparameter set consists of # of hidden dimensions [8, 16, 32, 64, 128], # of output dimension [8, 16, 32, 64, 128], learning rate [5e-5, 1e-4, 5e-4, 1e-3], batch size: [4, 8, 16, 32, 64] and dropout fraction [0.1, 0.3, 0.5, 0.7]. We use ELU as the activation function throughout. The optimal set for GCN is hidden dimension of size 8, output dimension of size 8, batch size 8, dropout rate 0.3 and learning rate 0.001. The optimal set of GraphSAGE is hidden dimension of size 8, output dimension of size 8, batch size 64, dropout rate 0.3 and learning rate 0.001. For GAT, the additional hyperparameter set includes number of attention dimension [8, 16, 32, 64, 128] and number of heads [2, 4, 8, 16, 32]. The optimal set is batch size 4, dropout rate 0.5, learning rate 0.001, output dimension 8, number of heads 32, number of attention head dimension 64. We optimize on the bond edge type and use the same set for KNN edge type. All models were trained on an internal cluster using a single NVIDIA V100 GPU.

### E Societal Impact

The potential risks of our library are minimal. A foreseeable hazardous application of our work is its use by nefarious actors to engineer harmful biomolecules, such as engineering toxins with greater efficacy. However, we believe these risks are minimal and shared across any developments in making computational design of biomolecules more accessible. We believe that utility of Graphein in the context of therapeutic development significantly outweighs these unlikely scenarios and anticipate our contribution to be a net force for social good and public health.

[7] Raji Heyrovska. Atomic structures of all the twenty essential amino acids and a tripeptide, with bond lengths as sums of atomic covalent radii, 2008.

[8] GetContacts. Getcontacts. URL https://getcontacts.github.io/.

[9] Jon C. Baber and Edward E. Hodgkin. Automatic assignment of chemical connectivity to organic molecules in the cambridge structural database. *Journal of Chemical Information and Modeling*, 32(5):401–406, September 1992. doi: 10.1021/ci00009a001. URL https://doi.org/10.1021/ci00009a001.

[10] Dong-Sheng Cao, Qing-Song Xu, and Yi-Zeng Liang. propy: a tool to generate various modes of Chou's PseAAC. *Bioinformatics*, 29(7):960–962, 02 2013. ISSN 1367-4803. doi: 10.1093/bioinformatics/btt072. URL https://doi.org/10.1093/bioinformatics/btt072.

[11] Thomas N Kipf and Max Welling. Semi-supervised classification with graph convolutional networks. *ICLR*, 2017.

[12] William L Hamilton, Rex Ying, and Jure Leskovec. Inductive representation learning on large graphs. In *NeurIPS*, pages 1025–1035, 2017.

[13] Petar Veličković, Guillem Cucurull, Arantxa Casanova, Adriana Romero, Pietro Lio, and Yoshua Bengio. Graph attention networks. *ICLR*, 2018.
